# PATZ1 remodels the nucleosome landscape to promote chromatin accessibility in injured neurons

**DOI:** 10.64898/2025.12.22.695949

**Authors:** Netra Krishna, Anisha S Menon, Manojkumar Kumaran, Dhruva Kumar Kesireddy, Ishwariya Venkatesh

**Affiliations:** Council of Scientific and Industrial Research (CSIR) – Centre for Cellular and Molecular Biology (CCMB), Hyderabad 500007, Telangana, India; Academy of Scientific and Innovative Research (AcSIR), Ghaziabad 201002, Uttar Pradesh, India

**Author notes:** These two authors contributed equally (Netra Krishna, Anisha S Menon).

## Abstract

Adult central nervous system neurons fail to regenerate after injury, in part due to epigenetic constraints that maintain a growth-restrictive chromatin state. We previously showed that the transcription factor PATZ1 reprograms chromatin accessibility in injured corticospinal neurons to promote axon regeneration, but the underlying mechanism remained unclear. Here we use nucleosome-resolution profiling to reveal that PATZ1 induces widespread nucleosome eviction at regeneration-associated gene loci. PATZ1 treatment dramatically reduces both nucleosome occupancy and fuzziness, indicating active chromatin remodeling rather than passive destabilization. This remodeling occurs preferentially at distal regulatory elements, where PATZ1 drives a greater than 3-fold expansion of H3K27ac-marked active enhancers. Nucleosome eviction at these sites precedes enhancer activation, establishing a mechanistic link between chromatin remodeling and transcriptional reprogramming. Our findings demonstrate that targeted nucleosome eviction at enhancers underlies PATZ1-mediated chromatin reorganization, providing mechanistic insight into how epigenetic barriers to CNS regeneration can be overcome.

## Introduction

Central nervous system neurons in adult mammals fail to regenerate following injury, a limitation underlying the permanent disability associated with spinal cord injury and neurodegenerative diseases^1–3^. While peripheral neurons retain regenerative capacity throughout life, CNS neurons progressively lose this ability during development^4^. Multiple factors contribute to this failure, including the inhibitory extracellular environment^5^ and cell-intrinsic barriers within the neurons themselves^1^. Among these intrinsic constraints, epigenetic mechanisms have emerged as fundamental regulators of regenerative competence^6–13,11,12^.The chromatin landscape of adult CNS neurons becomes increasingly restrictive, with developmental gene programs silenced through histone modifications and altered chromatin architecture. This epigenetic consolidation locks neurons into a mature, non-regenerative transcriptional state, suggesting that chromatin remodeling may be necessary to unlock the latent regenerative potential of injured CNS neurons.

We recently demonstrated that the transcription factor PATZ1 can reprogram the chromatin landscape of injured corticospinal neurons to reinstate a growth-permissive state, leading to re-opening of several growth relevant genes and enhancers^14^. PATZ1 treatment resulted in dramatic changes in chromatin accessibility and gene expression, activating regeneration-associated programs normally silenced in adult neurons. However, the precise molecular mechanisms by which PATZ1 remodels chromatin remain unclear. Chromatin accessibility assays provide a broad view of open versus closed regions but do not reveal the underlying nucleosome dynamics that govern regulatory element function. Nucleosomes, the fundamental units of chromatin packaging, can be characterized by two key parameters: occupancy, which reflects the probability that a nucleosome is positioned at a given locus, and fuzziness, which measures the precision of nucleosome positioning^15^. Nucleosome positioning at regulatory elements controls transcription factor access, and eviction from specific sites is required for factor binding and gene activation^16^. Examining how PATZ1 affects these properties can determine whether chromatin opening reflects active nucleosome remodeling or alternative mechanisms.

Here we present a comprehensive analysis of nucleosome dynamics following PATZ1 treatment in injured corticospinal neurons. We found that PATZ1 induces dramatic reductions in nucleosome occupancy at upregulated genes, accompanied by decreased fuzziness indicating active remodeling rather than random destabilization. Nucleosome eviction occurs preferentially at distal regulatory elements, where PATZ1 treatment results in a greater than 3-fold expansion of active enhancers marked by H3K27ac. These findings establish nucleosome eviction as the primary mechanism underlying PATZ1-mediated chromatin remodeling and identify enhancer activation as a key downstream consequence, providing mechanistic insight into how targeted chromatin remodeling can overcome the epigenetic barriers limiting CNS regeneration.

## Results

### Nucleosome organization remains stable in corticospinal neurons following spinal cord injury

Chromatin accessibility is fundamentally shaped by nucleosome positioning. To understand how spinal cord injury affects the chromatin landscape of corticospinal neurons at nucleosome resolution, and to establish a baseline against which therapeutic interventions could be assessed, we applied NucleoATAC ^15^ to our previous snATAC-seq datasets from L5 ET neurons ^14^ (**Figure. 1A,D**). This computational framework exploits the characteristic fragment size distribution of ATAC-seq data to infer nucleosome positions genome-wide. For each called nucleosome, NucleoATAC generates two complementary metrics: occupancy, representing the proportion of cells harbouring a nucleosome at that position (scale 0 to 1), and fuzziness, quantifying the positional variability of that nucleosome across the cell population (**Figure. 1B,C**). Together, these metrics capture both the prevalence and precision of nucleosome placement, features that directly influence transcription factor access and gene regulation.

**Figure 1.**
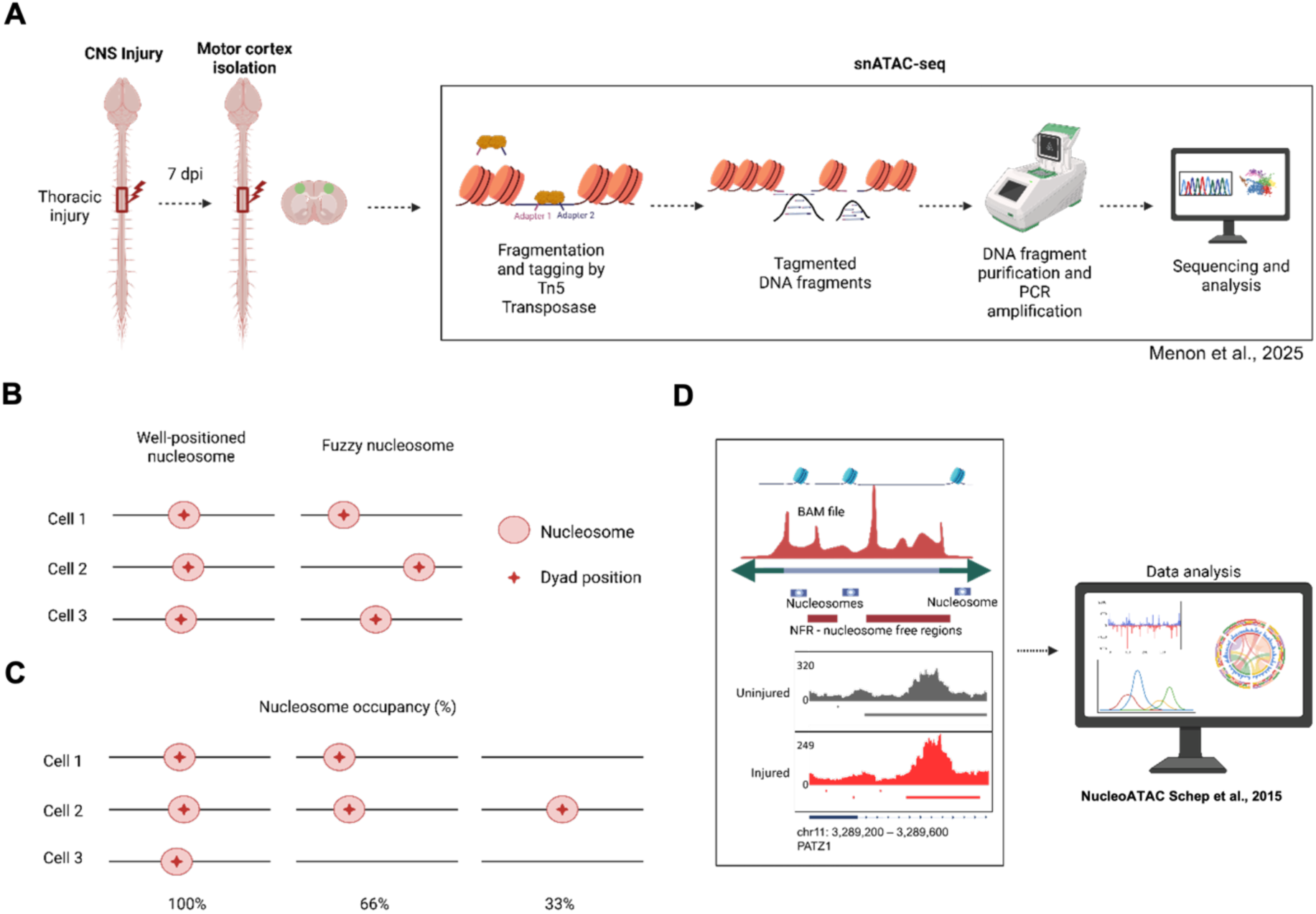
Analytical framework for nucleosome-level characterization of chromatin accessibility in corticospinal neurons. **(A)** Schematic of the single-nucleus ATAC-seq experimental design. Layer 5 extratelencephalic (L5 ET) neurons were isolated from mouse motor cortex under three conditions: uninjured, 1 week post-spinal cord injury (SCI), and SCI with AAV-mediated PATZ1 overexpression. Nuclei were processed for snATAC-seq to capture chromatin accessibility at single-cell resolution. **(B)** Conceptual illustration of nucleosome fuzziness. Fuzziness quantifies the positional consistency of a nucleosome across cells. A well-positioned nucleosome (left) exhibits low fuzziness, with consistent dyad placement across cells. A “fuzzy” nucleosome (right) shows high positional variability, indicating heterogeneous chromatin organization across the cell population. **(C)** Conceptual illustration of nucleosome occupancy. Occupancy represents the fraction of cells in which a given genomic position is occupied by a nucleosome. High occupancy (left, 100%) indicates consistent nucleosome placement; intermediate occupancy (middle, 66%) reflects partial population heterogeneity; low occupancy (right, 33%) suggests the position is nucleosome-depleted in most cells. **(D)** Computational workflow for nucleosome calling using NucleoATAC. ATAC-seq fragments corresponding to mono-nucleosomal sizes (∼147 bp ± linker) were analyzed using NucleoATAC (Schep et al., 2015) to infer nucleosome positions, occupancy scores (0–1 scale), and fuzziness values genome-wide.

To assess whether spinal cord injury alters nucleosome organization in corticospinal neurons, we first examined the quality of nucleosome signals in our snATAC-seq data. V-plots displaying ATAC-seq fragment sizes relative to inferred nucleosome dyad positions revealed the expected “V” pattern in both uninjured and injured conditions, with fragment sizes centered around 147 bp, confirming robust detection of canonical nucleosomal DNA protection (**Figure.2A,B, Supplementary table S2,3**).

Direct comparison of occupancy distributions revealed nearly identical central tendencies and dispersions between conditions (**Figure. 2C**). Median occupancy was 0.105 in uninjured neurons and 0.100 in injured neurons, a difference of 4.8%. Mean occupancy values were similarly comparable (0.124 vs. 0.121). Although statistical testing yielded significant p-values (Dunn Kruskal-Wallis multiple comparison adjusted with Bonferroni method Z =-18.97833, p = 7.73 × 10⁻⁸⁰), the effect size was negligible (Cliff’s δ = 0.046), indicating that the statistical significance reflects the large sample sizes rather than biologically meaningful differences. Interquartile ranges were also comparable between conditions (IQR: 0.099 uninjured vs. 0.107 injured).

**Figure 2.**
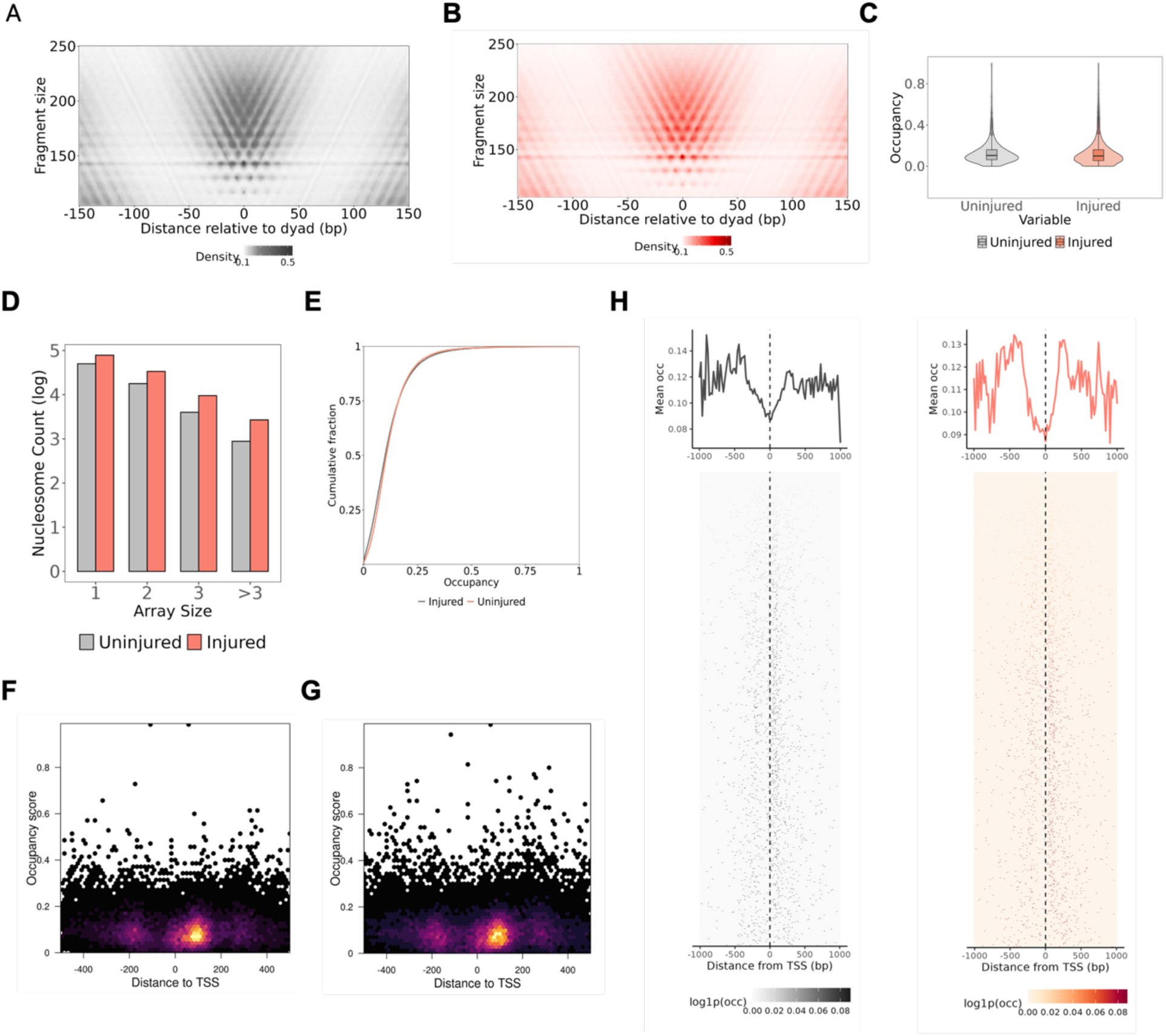
Nucleosome occupancy is unchanged between uninjured and injured corticospinal neurons. **(A)** V-plot showing ATAC-seq fragment size distribution relative to nucleosome dyad positions in uninjured L5 ET neurons (n = 72,576 nucleosome calls). The characteristic “V” pattern reflects protection of nucleosomal DNA, with fragment sizes clustering around 147 bp at the dyad center. **(B)** V-plot for injured (1 week post-SCI) L5 ET neurons (n = 123,950 nucleosome calls) displaying a similar fragment size distribution pattern to uninjured controls. **(C)** Violin plots with embedded box plots comparing nucleosome occupancy distributions. Median occupancy: uninjured = 0.105, injured = 0.100. Despite significant p-values due to large sample sizes (Dunn Kruskal-Wallis multiple comparison adjusted with Bonferroni method Z =-18.97833, p = 7.73 × 10⁻⁸⁰), effect size was negligible (Cliff’s delta = 0.046). **(D)** Nucleosome array size distribution comparing uninjured (gray) and injured (red) conditions. Array sizes represent the number of consecutive, regularly spaced nucleosomes detected within 250bp of each other (log scale on y-axis). **(E)** Empirical cumulative distribution function (ECDF) of nucleosome occupancy scores showing near-complete overlap between uninjured (gray) and injured (red) distributions. **(F)** Hexbin density plot of nucleosome occupancy as a function of distance to the nearest transcription start site (TSS) in uninjured neurons. **(G)** Hexbin density plot of occupancy versus TSS distance for injured neurons, showing a comparable spatial distribution to uninjured controls. **(H)** Heatmap plots indicating the nucleosome positions and their log occupancy values 1kb upstream and downstream of the TSS. The left panel is uninjured control and the right panel is the injured condition.

We next compared nucleosome organization between conditions across multiple metrics. Nucleosome array distributions, reflecting the periodic spacing of consecutive nucleosomes along chromatin fibers, were comparable between uninjured and injured neurons across mono-, di-, tri-, and higher-order arrays (**Figure. 2D, Supplementary table S5,6**). Empirical Cumulative Distribution Function (ECDF) analysis, plot analysis revealed that occupancy cumulative distributions were nearly superimposed across the full range of occupancy scores (**Figure. 2E, Supplementary table S5,6**).

Finally, we examined the spatial relationship between nucleosome occupancy and genomic features. Hexbin density plots of occupancy versus distance to the nearest TSS revealed similar distributions in both conditions, with nucleosomes present across promoter-proximal and distal regions at comparable occupancy levels (**Figure. 2F,G, Supplementary table S5,6**). This comparable mean occupancy is also seen in the proximity of the TSS within 1kb (**Figure 2H, top panel, Supplementary table 5,6**), as well as similar count and positioning of nucleosome calls around the TSS (**Figure 2H, bottom panel, Supplementary table 5,6**).

Together, these analyses demonstrate that spinal cord injury does not substantially alter nucleosome occupancy in L5 ET neurons. This stability is consistent with prior observations that central nervous system neurons fail to mount robust transcriptional or epigenetic responses to thoracic injury at the cell body level in CST neurons^14,17^.

### Nucleosome positioning precision is maintained following spinal cord injury

Having established that nucleosome occupancy is unchanged following injury, we next examined whether spinal cord injury affects nucleosome positioning precision. Fuzziness quantifies the variability in nucleosome position across cells, with high values indicating heterogeneous placement and low values indicating consistent positioning across the cell population^15^.

Direct comparison of distributions revealed nearly identical central tendencies (**Figure. 3A,B**). Overlaid histograms demonstrated similar distribution shapes with peak density around fuzziness values of 20-25 (**Figure. 3A, Supplementary table S5,6**). Median fuzziness was 23.55 in uninjured neurons (n = 72,576) and 22.81 in injured neurons (n = 123,950), a difference of 3.1% (**Figure. 3B, Supplementary table S5,6**). Although statistical testing yielded significant p-values (Wilcoxon rank sum test with continuity correction= 6.97 × 10⁹, p = 2.2 × 10⁻¹⁶), the effect size was negligible (Cliff’s δ = 0.092), confirming that the statistical significance reflects large sample sizes rather than biologically meaningful differences.

**Figure 3.**
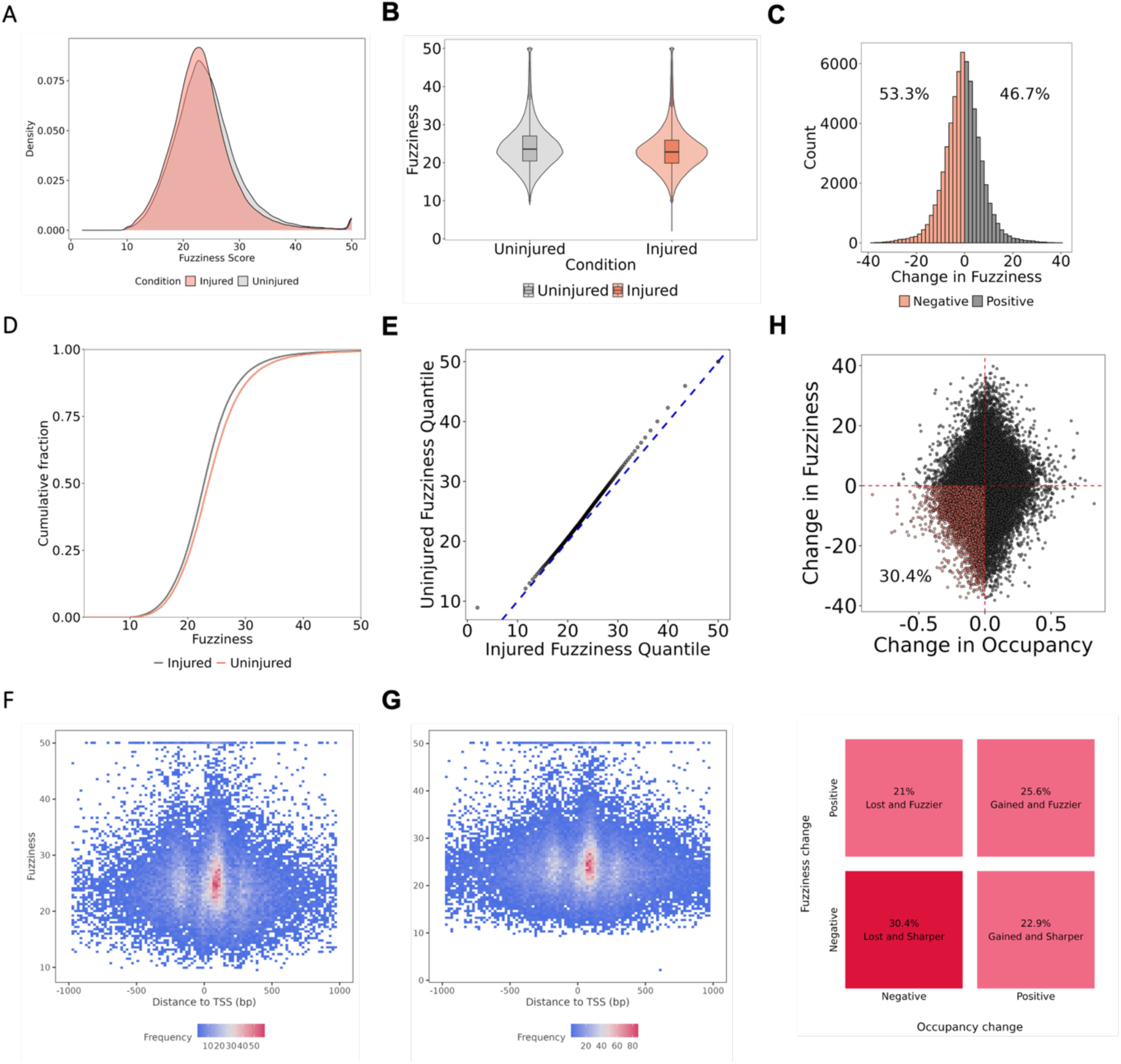
Nucleosome positioning precision (fuzziness) is unchanged between uninjured and injured corticospinal neurons. **(A)** Overlaid density histograms of fuzziness values for uninjured (gray) and injured (red) conditions, demonstrating similar distribution shapes with peaks around 20-25. **(B)** Violin plots with embedded box plots comparing fuzziness distributions. Median fuzziness: uninjured = 23.55, injured = 22.81 (3.1% difference). Despite significant p-values due to large sample sizes (Wilcoxon rank sum test with continuity correction= 6.97 × 10⁹, p = 2.2 × 10⁻¹⁶), effect size was negligible (Cliff’s delta = 0.092). **(C)** Histogram of change in fuzziness (Δ fuzziness = injured minus uninjured) for nucleosomes detected in both conditions. Red bars indicate nucleosomes with decreased fuzziness following injury while gray bars indicate increased fuzziness. **(D)** Empirical cumulative distribution function (ECDF) of nucleosome fuzziness showing near-complete overlap between uninjured (gray) and injured (red) distributions. **(E)** Quantile-quantile (Q-Q) plot comparing fuzziness distributions between uninjured (x-axis) and injured (y-axis) conditions. Points closely follow the identity line (blue dashed), indicating concordant distributions (Kolmogorov-Smirnov D = 0.074). **(F)** 2D density plot of nucleosome fuzziness versus distance to nearest TSS in uninjured neurons. **(G)** 2D density plot of fuzziness versus TSS distance for injured neurons, showing comparable spatial distribution to uninjured controls. **(H)** Cin occupancy (Δ occupancy) versus change in fuzziness (Δ fuzziness) for matched nucleosomes. The above panel is a scatterplot describing the data. Red points indicate nucleosomes with decreased fuzziness; black points indicate increased fuzziness. The weak correlation (Kendall tau = 0.1) suggests that changes in occupancy and fuzziness are largely independent. The lower panel highlights the percentage distribution across all four quadrants

To directly assess position-specific changes, we identified nucleosomes detected at the same genomic coordinates in both uninjured and injured conditions (n = 1,458 matched nucleosomes) and calculated the change in fuzziness for each. The distribution of Δ fuzziness was centred near zero (median =-0.39, mean =-0.75), with 53.3% of nucleosomes showing decreased fuzziness and 46.7% showing increased fuzziness following injury (**Figure. 3C, Supplementary table S5,6**). Although the Wilcoxon signed-rank test indicated a statistically significant shift (p = 3.47 × 10⁻⁷), the magnitude of this change was minimal relative to the overall fuzziness range.

Genome-wide comparison of fuzziness distributions confirmed the positioning stability. ECDF curves were nearly superimposed across the full range of fuzziness values (**Figure. 3D, Supplementary table S5,6**). and Q-Q plot analysis revealed that fuzziness quantiles from injured neurons tracked closely along the identity line relative to uninjured controls (**Figure. 3E, Supplementary table S5,6**).

The spatial relationship between fuzziness and genomic features was also preserved. 2-dimensional density plots of fuzziness versus distance to TSS revealed similar patterns in both conditions, with nucleosomes showing consistent fuzziness values across promoter-proximal and distal regions (**Figure. 3F,G, Supplementary table S5,6**).

Finally, we examined whether changes in occupancy and fuzziness were correlated at individual nucleosome positions. Scatterplot analysis of Δ occupancy versus Δ fuzziness revealed only a weak positive correlation (Kendall τ = 0.1, p = 2.2 × 10⁻¹⁶), indicating that these two metrics change largely independently (**Figure. 3H**). Nucleosomes with decreased occupancy did not systematically show altered positioning precision, and vice versa.

Together with the occupancy analyses in Figure 2, these results demonstrate that spinal cord injury does not substantially alter nucleosome organization in L5 ET neurons. Both the prevalence of nucleosome placement (occupancy) and the precision of positioning (fuzziness) remain stable following axonal disconnection. This chromatin stability is consistent with the failure of central nervous system neurons to initiate regenerative transcriptional programs following injury, and establishes the baseline against which active chromatin remodeling interventions can be assessed.

### PATZ1 dramatically reduces nucleosome occupancy in injured corticospinal neurons

Having established that spinal cord injury alone does not alter nucleosome organization in L5 ET neurons (**Figures 2-3**), we next examined whether PATZ1 overexpression could actively remodel the chromatin landscape. We recently demonstrated that PATZ1 primes regeneration-specific loci in injured neurons by increasing chromatin accessibility at growth-associated genes (Menon et al., 2025). We hypothesized that this activity would manifest as altered nucleosome properties genome-wide.

V-plot analysis revealed qualitative differences between injured and PATZ1-treated conditions (**Figure. 4A,B**). While both showed the characteristic “V” pattern indicative of nucleosomal protection, signal intensity was notably attenuated in PATZ1-treated neurons, suggesting reduced nucleosome occupancy. Direct comparison of occupancy distributions revealed an 86% reduction in median occupancy following PATZ1 treatment (0.100 injured vs. 0.014 PATZ1) (**Figure. 4C,Supplementary table S6,7**). This difference was highly significant (Dunn Kruskal-Wallis multiple comparison adjusted with Bonferroni method indicated a Z score = 259.492), with a medium effect size (Cliff’s δ = 0.603). The magnitude of this change stands in stark contrast to the negligible differences observed between uninjured and injured conditions (Cohen’s d = 0.03).

**Figure 4.**
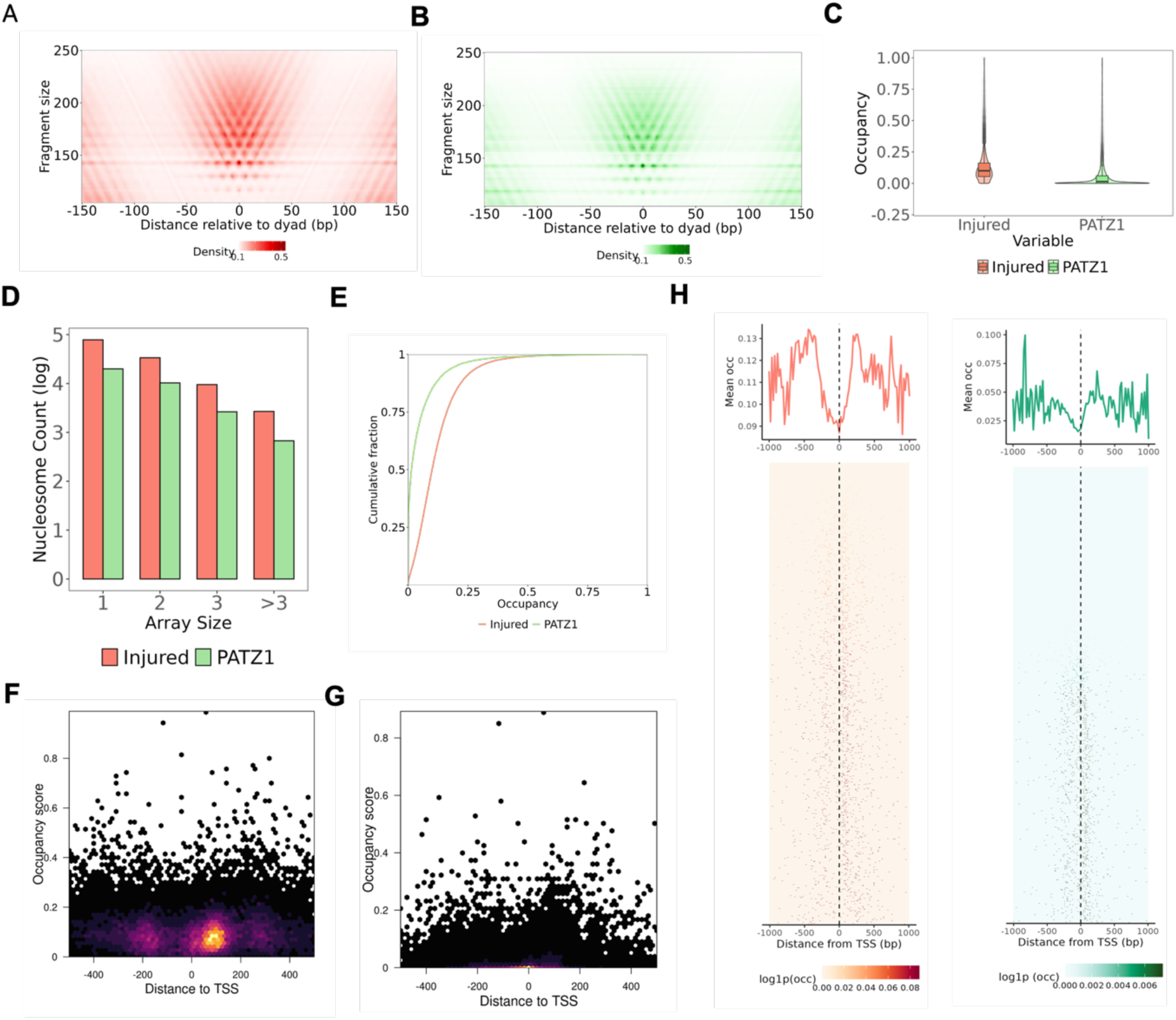
PATZ1 introduction results in global occupancy reduction in injured corticospinal neurons. **(A)** V-plot showing ATAC-seq fragment size distribution relative to nucleosome dyad positions in injured L5 ET neurons (n = 123,950 nucleosome calls). The characteristic “V” pattern reflects protection of nucleosomal DNA, with fragment sizes clustering around 147 bp at the dyad center. **(B)** V-plot for PATZ1 treated injured (1 week post-SCI) L5 ET neurons (n = 33,448 nucleosome calls) displaying a similar fragment size distribution pattern to the injured controls as expected. **(C)** Violin plots with embedded box plots comparing nucleosome occupancy distributions. Median occupancy: injured = 0.100 and PATZ1 = 0.013. Dunn Kruskal-Wallis multiple comparison adjusted with Bonferroni method indicated a Z score = 259.492 and effect size was found to be large (Cliff’s delta = 0.603) **(D)** Nucleosome array size distribution comparing injured (red) and PATZ1 (green) conditions. Array sizes represent the number of consecutive, regularly spaced nucleosomes detected within 250bp of each other (log scale on y-axis). **(E)** Empirical cumulative distribution function (ECDF) of nucleosome occupancy scores showing higher distribution around lower occupancy in PATZ1 (green) compared to injured (red) condition. **(F)** Hexbin density plot of nucleosome occupancy as a function of distance to the nearest transcription start site (TSS) in injured neurons. **(G)** Hexbin density plot of occupancy versus TSS distance for PATZ1 treated injured neurons, showing a reduced occupancy distribution compared to injured controls. **(H)** Heatmap-lineplots indicating the nucleosome positions and their log occupancy values 1kb upstream and downstream of the TSS. The left panel is injured condition and the right panel is the PATZ1 treated injured condition showing overall reduced nucleosome calls within 1kb of the TSS.

PATZ1 treatment also resulted in fewer total nucleosomes called (33,448 vs. 123,950 in injured), with reductions evident across all array sizes (**Figure. 4D, Supplementary table S3,4**). Higher-order arrays (>3 consecutive nucleosomes) showed the largest proportional decrease, suggesting that PATZ1 may preferentially disrupt regularly spaced nucleosome arrays.

Quantitative analysis confirmed that PATZ1 induces a dramatic reduction in nucleosome occupancy. ECDF curves demonstrated a pronounced leftward shift, with 50% of PATZ1 nucleosomes falling below an occupancy of 0.014 compared to 0.100 for injured controls (**Figure. 4E, Supplementary table S6,7**).

The spatial relationship between occupancy and genomic features was also altered by PATZ1 treatment. Hexbin plots of occupancy versus distance to TSS revealed that while injured neurons showed nucleosomes distributed across a range of occupancy values at all distances (**Figure. 4F**), PATZ1 treatment compressed this distribution toward lower occupancy values genome-wide (**Figure. 4G**, **Supplementary table S6,7**). The reduced occupancy induced by PATZ1 is also seen in the proximity of the TSS within 1kb (**Figure 4H, top panel, Supplementary table 6,7**), as well as depleted nucleosome calls around the TSS across all the genes (**Figure 4H, bottom panel, Supplementary table 6,7**). This suggests that PATZ1-mediated nucleosome remodeling is not restricted to specific genomic contexts but rather reflects a global shift in chromatin organization, consistent with its role in priming regeneration-associated loci for transcriptional activation.

### PATZ1 increases nucleosome positioning precision in injured corticospinal neurons

In addition to reducing nucleosome occupancy (**Figure 4**), we examined whether PATZ1 affects nucleosome positioning precision as measured by fuzziness. Fuzziness quantifies the variability in nucleosome position across cells; lower fuzziness indicates more consistent nucleosome placement within the cell population.

Direct comparison of fuzziness distributions revealed a 23% reduction in median fuzziness following PATZ1 treatment (22.81 injured vs. 17.64 PATZ1) (**Figure. 5A,B, Supplementary table S6,7**). This difference was highly significant (Dunn Kruskal-Wallis multiple comparison adjusted with Bonferroni method Z = 155.9459, p = 0) with a large effect size (Cliff’s δ = 0.57). Notably, the effect size for fuzziness exceeded that for occupancy (Cliff’s δ = 0.092), indicating that PATZ1 has an even more pronounced effect on positioning precision than on occupancy per se.

**Figure 5.**
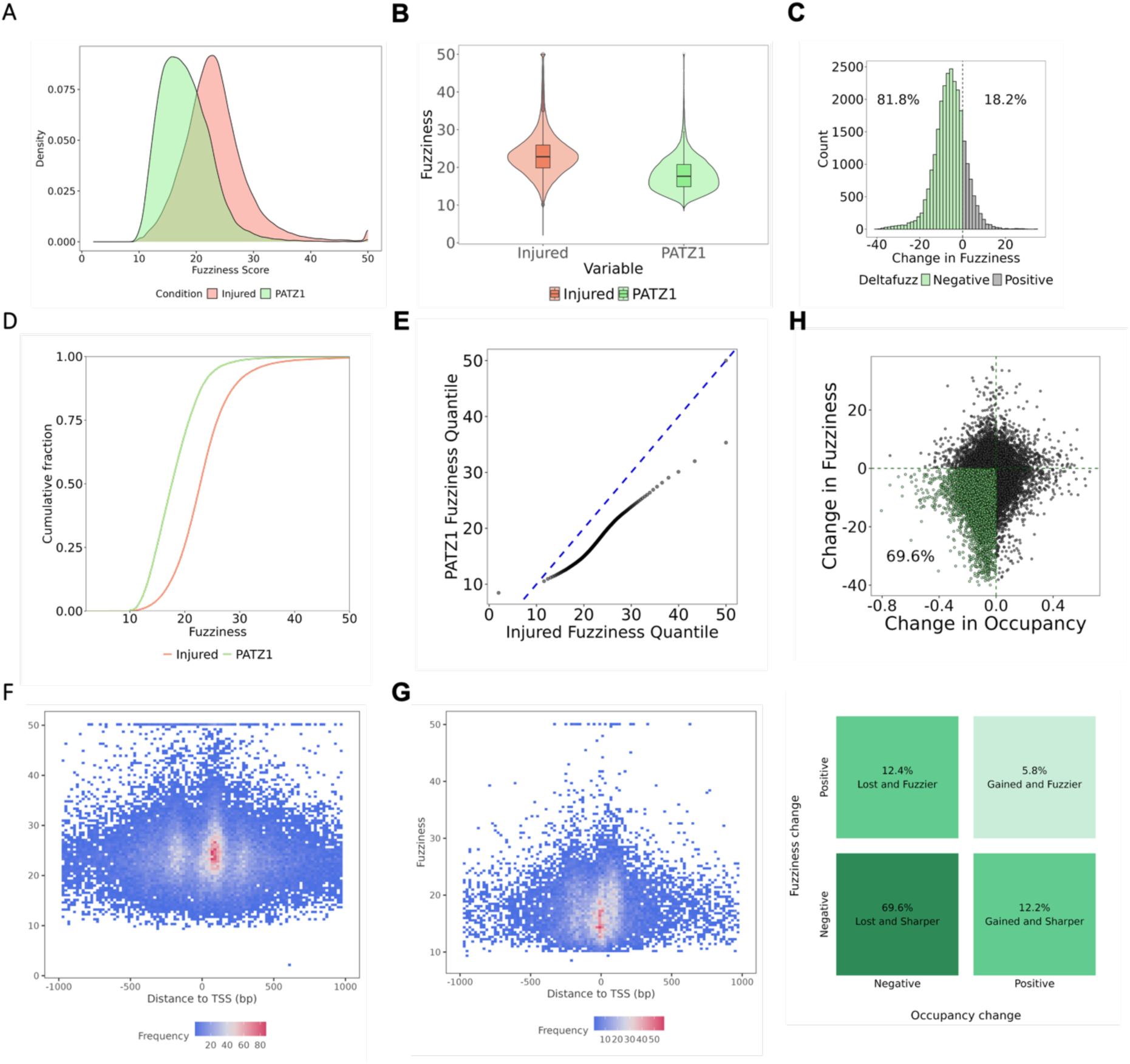
Introduction of PATZ1 to injured corticospinal neurons results in more precise nucleosome positioning. **(A)** Overlaid density histograms of fuzziness values for and injured (red) and PATZ1 (green) conditions, wherein PATZ1 distribution peak falls between 12-15, far lower than the injured condition. **(B)** Violin plots with embedded box plots comparing fuzziness distributions. Median fuzziness: injured = 22.81, PATZ1 = 17.63 (3.1% difference. **(C)** Histogram of change in fuzziness (Δ fuzziness = injured minus PATZ1) for nucleosomes detected in both conditions. Green bars indicate nucleosomes with decreased fuzziness following PATZ1 treatment; gray bars indicate increased fuzziness. **(D)** Empirical cumulative distribution function (ECDF) indicates a higher fraction of low fuzziness scores in PATZ1 (green) compared to the injured (red) ditrsibution. **(E)** Quantile-quantile (Q-Q) plot comparing fuzziness distributions between injured (x-axis) and PATZ1 treatment (y-axis) conditions. Points deviate from the identity line (blue dashed; Kolmogorov-Smirnov D = 0.074). **(F)** 2D density plot of nucleosome fuzziness versus distance to nearest TSS in injured neurons. **(G)** 2D density plot of fuzziness versus TSS distance for PATZ1 treated injured neurons, showing shifted spatial distribution in PATZ1 compared to injured controls. **(H)** Change in occupancy (Δ occupancy) versus change in fuzziness (Δ fuzziness) for matched nucleosomes. The above panel is a scatterplot describing the data. Green points indicate nucleosomes with decreased fuzziness; black points indicate increased fuzziness. The weak correlation (Kendall tau = 0.14) suggests that changes in occupancy and fuzziness are largely independent. The lower panel highlights the percentage distribution across all four quadrants

To directly assess position-specific changes, we first analyzed nucleosomes detected at the same genomic coordinates in both conditions (n = 554 matched positions). The distribution of Δ fuzziness was strongly skewed toward negative values, with 84.8% of matched nucleosomes showing decreased fuzziness following PATZ1 treatment (**Figure. 5C, Supplementary table S6,7**). The median reduction was 4.44 units (Wilcoxon signed-rank p = 1.79 × 10⁻⁷¹), representing a substantial shift toward more precise positioning.

ECDF curves demonstrated a pronounced leftward shift in the PATZ1 distribution (**Figure. 5D, top, Supplementary table S6,7**). Q-Q plot analysis confirmed that PATZ1 fuzziness quantiles fell systematically below those of injured neurons across the entire distribution (**Figure. 5E, Supplementary table S6,7**). Unlike the uninjured versus injured comparison where points tracked the identity line (**Figure 3B**), PATZ1 treatment produced consistent deviation below the line, indicating reduced fuzziness at all quantiles.

Examination of the spatial distribution of fuzziness revealed important insights into PATZ1 function. In injured neurons, fuzziness values clustered around 20-25 across both promoter-proximal and distal regions (**Figure. 5E, Supplementary table S6,7**). Following PATZ1 treatment, fuzziness shifted toward lower values (clustering around 15-20) with pronounced effects evident at TSS-distal regions (**Figure. 5F, Supplementary table S6,7**). This pattern is consistent with PATZ1 promoting chromatin accessibility at distal regulatory elements, including enhancers that control regeneration-associated gene expression^14^. The increased positioning precision at these regions would facilitate transcription factor binding and enhancer-promoter communication necessary for activating regenerative transcriptional programs.

Finally, we examined whether changes in occupancy and fuzziness were correlated at individual nucleosome positions. Scatterplot analysis revealed only a weak positive correlation (Kendall τ = 0.21, p = 2.2 × 10⁻^16^), indicating that these two metrics change largely independently (**Figure. 5G, Supplementary table S6,7**). However, the majority of nucleosomes clustered in the lower-left quadrant (decreased occupancy, decreased fuzziness), consistent with coordinated chromatin remodeling that both reduces nucleosome barrier function and increases positioning precision.

Together with the occupancy findings (Figure 4), these results demonstrate that PATZ1 induces a coordinated reorganization of nucleosome properties in injured corticospinal neurons. PATZ1 not only reduces the fraction of cells with nucleosomes at given positions (occupancy) but also increases the consistency of nucleosome placement among the remaining nucleosomes (reduced fuzziness). This dual effect, particularly prominent at distal regulatory regions, is consistent with active chromatin remodeling that opens enhancer elements and creates a permissive chromatin environment for regeneration-associated gene activation, as we recently demonstrated ^14^.

### PATZ1 Induces Nucleosome Depletion at Differentially Accessible Chromatin Regions

To understand the chromatin remodeling mechanisms underlying PATZ1-mediated accessibility changes, we performed nucleosome-level analysis of differentially accessible snATAC-seq peaks using NucleoATAC. Integration of nucleosome calls with PATZ1-responsive peaks revealed substantial nucleosome depletion following PATZ1 treatment. Of the differentially accessible peaks, 81.3% lacked detectable nucleosomes in the PATZ1 condition, while 11.1% showed nucleosome eviction—peaks that contained nucleosomes in injured neurons but lost them upon PATZ1 treatment (**Figure. 6A, Supplementary table S8**). An additional 4.6% retained nucleosomes and 3.1% showed nucleosome appearance in both conditions.

**Figure 6.**
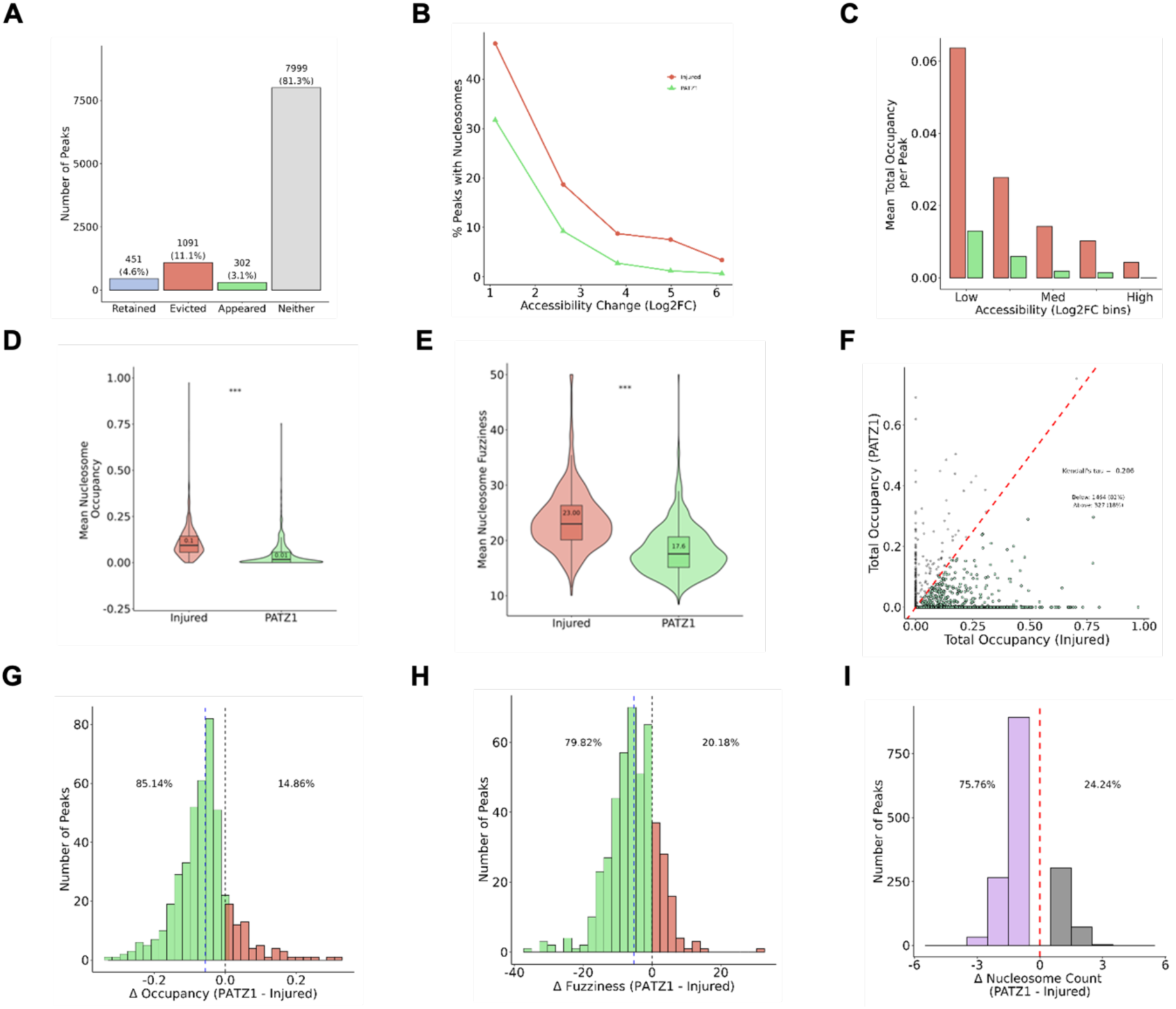
Integration of differentially accessible snATAC-seq peaks reveals depleted occupancy in PATZ1 nucleosome calls. (A) Bar plot showing the presence and/or absence of nucleosomes in PATZ1 differentially accessible peaks. 11.1% of the peaks show evicted nucleosomes in the PATZ1 condition. (B) Line plot showing the decreasing trend of nucleosome peak percentage with the log2 fold change in accessibility. (C) Bar plot showing the mean nucleosome occupancy per peak against low, medium, and high differentially accessible regions. (D) Violin boxplots comparing the mean nucleosome occupancy in differentially accessible peaks between injured (red) neurons and PATZ1-treated (green) injured neurons. Median occupancy injured – 0.1; PATZ1 – 0.01 (E) Violin boxplots comparing the mean nucleosome fuzziness in differentially accessible peaks between injured (red) neurons and PATZ1-treated (green) injured neurons. Median fuzziness injured – 23.00; PATZ1 – 17.58 (F) Scatterplot comparing matched nucleosome call occupancy between injured and PATZ1 conditions. Kendall’s tau was calculated with a tau = 0.206 indicating independence in occupancy between injured and PATZ1. (G) Histogram highlighting the number of differentially accessible peaks against occupancy change (Δ occupancy) for PATZ1 and injured matched nucleosomes. 85% of the peaks show reduced occupancy compared to injured neurons. (H) Histogram highlighting the number of differentially accessible peaks against fuzziness change (Δ fuzziness) for PATZ1 and injured matched nucleosomes. 80% of the peaks show reduced occupancy compared to injured neurons. (I) Discrete histogram showing the difference in number of nucleosomes between injured and PATZ1 conditions with the associated number of peaks. 75% of the peaks are nucleosome-evicted (purple) in the case of PATZ1-treated injured neurons

The relationship between accessibility gain and nucleosome presence followed a clear inverse trend: the percentage of peaks containing nucleosomes decreased progressively with increasing accessibility fold change, dropping from approximately 40% at low fold changes to below 10% at the highest accessibility gains (**Figure. 6B, Supplementary table S8**). This pattern was consistent across both injured and PATZ1 conditions, though PATZ1-treated neurons showed consistently lower nucleosome occupancy across all accessibility bins. Correspondingly, mean total nucleosome occupancy per peak was markedly reduced in high-accessibility regions compared to low and medium accessibility bins (**Figure. 6C**).

Direct comparison of nucleosome occupancy between conditions demonstrated that PATZ1 treatment significantly reduced mean nucleosome occupancy at differentially accessible peaks (median occupancy: injured = 0.1, PATZ1 = 0.01; p < 0.001; **Figure. 6D, Supplementary table S8**). PATZ1 treatment also significantly decreased nucleosome fuzziness, a measure of nucleosome positioning variability, indicating more precise positioning of remaining nucleosomes (median fuzziness: injured = 23.00, PATZ1 = 17.58; p < 0.001; **Figure. 6E, Supplementary table S8**).

To assess whether occupancy changes were coordinated across conditions, we compared matched nucleosome calls between injured and PATZ1-treated neurons. Scatterplot analysis revealed weak correlation between occupancy values (Kendall’s τ = 0.206), suggesting that PATZ1 induces condition-specific rather than proportional changes in nucleosome positioning (**Figure. 6F, Supplementary table S8**). Examining the distribution of occupancy changes (Δ occupancy) showed that 85% of differentially accessible peaks exhibited reduced nucleosome occupancy following PATZ1 treatment compared to injured controls (**Figure. 6G**). Similarly, 80% of peaks showed reduced nucleosome fuzziness in PATZ1-treated neurons (**Figure. 6H, Supplementary table S8**). Analysis of nucleosome count differences revealed that 75.76% of peaks lost nucleosomes in the PATZ1 condition, while only 24.24% gained nucleosomes (**Figure. 6I, Supplementary table S8**).

Together, these findings demonstrate that PATZ1 promotes chromatin accessibility through active nucleosome eviction and depletion, establishing a permissive chromatin environment at regeneration-associated loci.

### PATZ1-Mediated Nucleosome Remodeling at Differentially Expressed Genes Reveals Pathway-Specific Chromatin Reorganization

To link nucleosome dynamics with transcriptional outcomes, we integrated snRNA-seq differentially expressed genes with nucleosome positioning data from accessible gene regions. Analysis of nucleosome fate at these loci revealed that the majority of regions lacked detectable nucleosomes, while 20% of nucleosomes present in injured neurons were evicted upon PATZ1 introduction (**Figure. 7A, Supplementary table S9**). A smaller fraction showed retained nucleosomes or nucleosome appearance in the PATZ1 condition.

**Figure 7.**
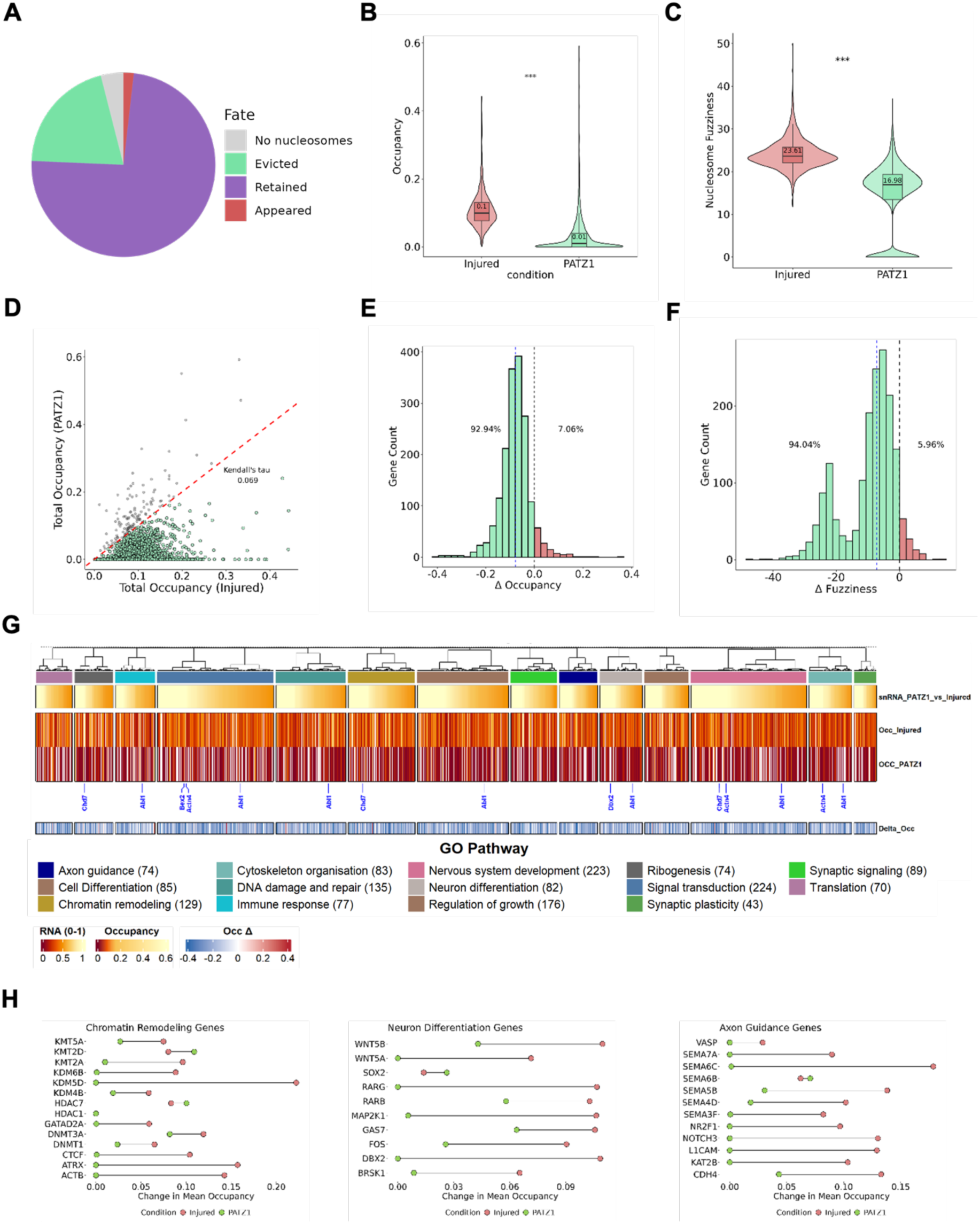
Integration of snRNA-seq differentially expressed genes reveals depleted occupancy and sharper positioning of nucleosomes in PATZ1 condition. **(A)** Pie chart depicting the nucleosome fate distribution in differentially expressed genes. 20% of the nucleosomes are evicted upon introduction of PATZ1 to injured neurons **(B)** Violin boxplots comparing the mean nucleosome occupancy in differentially accessible gene regions between injured (red) neurons and PATZ1-treated (green) injured neurons. Median occupancy injured – 0.1; PATZ1 – 0.01. **(C)** Violin boxplots comparing the mean nucleosome fuzziness in differentially accessible peaks between injured (red) neurons and PATZ1-treated (green) injured neurons. Median fuzziness injured – 23.61; PATZ1 – 16.98 (D) Scatterplot comparing matched nucleosome call occupancy between injured and PATZ1 conditions. Kendall’s tau was calculated: tau = 0.069 indicating independence in occupancy between injured and PATZ1. **(E)** Histogram highlighting the number of differentially expressed genes against occupancy change (Δ occupancy) for PATZ1 and injured matched nucleosomes. 93% of the peaks show reduced occupancy compared to injured neurons. **(F)** Histogram highlighting the number of differentially expressed genes against fuzziness change (Δ fuzziness) for PATZ1 and injured matched nucleosomes. 94% of the peaks show reduced occupancy compared to injured neurons. **(G)** Heatmap of multiple GO pathways comparing differentially expressed genes of from snRNA-seq, with occupancy scores from injured neurons and PATZ1-treated injured neurons, as well as the occupancy change score. **(H)** Lollipop plots showing the occupancy change across canonical genes across three different GO pathways.

Consistent with observations at differentially accessible peaks, PATZ1 treatment significantly reduced mean nucleosome occupancy at gene-associated accessible regions (median occupancy: injured = 0.1, PATZ1 = 0.01; p < 0.001; Fig. 7B). Nucleosome fuzziness was also significantly decreased following PATZ1 treatment, indicating sharper nucleosome positioning at remaining occupied sites (median fuzziness: injured = 23.61, PATZ1 = 16.98; p < 0.001; **Figure. 7C, Supplementary table S9**). Comparison of matched nucleosome calls between conditions revealed minimal correlation (Kendall’s τ = 0.069), suggesting that PATZ1 induces largely independent chromatin remodeling rather than proportional scaling of pre-existing nucleosome architecture (**Figure. 7D, Supplementary table S9**).

The directionality of nucleosome changes was strikingly consistent across differentially expressed genes. Analysis of occupancy differences showed that 92.94% of genes exhibited reduced nucleosome occupancy following PATZ1 treatment, with only 7.06% showing increased occupancy (**Figure. 7E, Supplementary table S9**). Similarly, 94.04% of genes displayed reduced nucleosome fuzziness in PATZ1-treated neurons compared to 5.96% with increased fuzziness (**Figure. 7F, Supplementary table S9**).

To determine whether nucleosome remodeling occurred preferentially at specific functional gene classes, we performed pathway-level analysis of occupancy changes. Heatmap visualization of GO pathway-annotated genes revealed coordinated nucleosome depletion across diverse functional categories, including nervous system development (223 genes), chromatin remodeling (129 genes), DNA damage and repair (135 genes), signal transduction (224 genes), and regulation of growth (176 genes) (**Figure. 7G, Supplementary table S9**). Genes within these pathways showed consistently higher occupancy in injured neurons compared to PATZ1-treated neurons, with negative occupancy change scores reflecting widespread nucleosome eviction. Notably, axon guidance (74 genes), synaptic signaling (89 genes), and synaptic plasticity (43 genes) pathways—directly relevant to regenerative capacity—displayed pronounced occupancy reduction.

Examination of individual genes within key pathways confirmed these patterns (**Figure. 7H, Supplementary table S9**). Chromatin remodeling genes including histone methyltransferases (*KMT5A*, *KMT2D*), demethylases (*KDM6A*, *KDM6B*, *KDM5D*, *KDM4B*), deacetylases (*HDAC7*, *HDAC1*), and DNA methyltransferases (*DNMT3A*, *DNMT3B*) all showed substantial reductions in nucleosome occupancy upon PATZ1 treatment. Neuron differentiation genes, including *WNT5B*, *WNT5A*, *SOX2*, *RARG*, *RARB*, *MAP2K1*, *GAS7*, *FOS*, and *DBX2*, similarly demonstrated occupancy decreases. Axon guidance genes critical for regeneration—including *VASP*, multiple semaphorins (*SEMA7A*, *SEMA6C*, *SEMA6B*, *SEMA6D*, *SEMA4D*, *SEMA3F*), *PAK1*, *NOTCH3*, *L1CAM*, *KIF2B*, and *CDH4*—exhibited reduced nucleosome occupancy in PATZ1-treated neurons.

These findings demonstrate that PATZ1 induces coordinated nucleosome depletion across functionally coherent gene programs, establishing a permissive chromatin landscape at loci governing neuronal differentiation, axon guidance, and synaptic function that collectively support the regenerative response.

### PATZ1 Drives Nucleosome Eviction at Enhancer Regions Including Novel PATZ1-Specific Regulatory Elements

To examine whether PATZ1-mediated chromatin remodeling extends to distal regulatory elements, we analyzed nucleosome dynamics at enhancer regions marked by H3K27ac. Classification of enhancer regions revealed four distinct categories based on their chromatin state: injured-specific enhancers (1134), PATZ1-only enhancers (2842), shared enhancers present in both conditions (3735), and PATZ1-specific enhancers uniquely accessible following treatment (893) (**Figure. 8A, Supplementary table S10**). Notably, PATZ1 treatment opened numerous enhancer regions not accessible in injured neurons, representing a substantial expansion of the active enhancer landscape.

**Figure 8.**
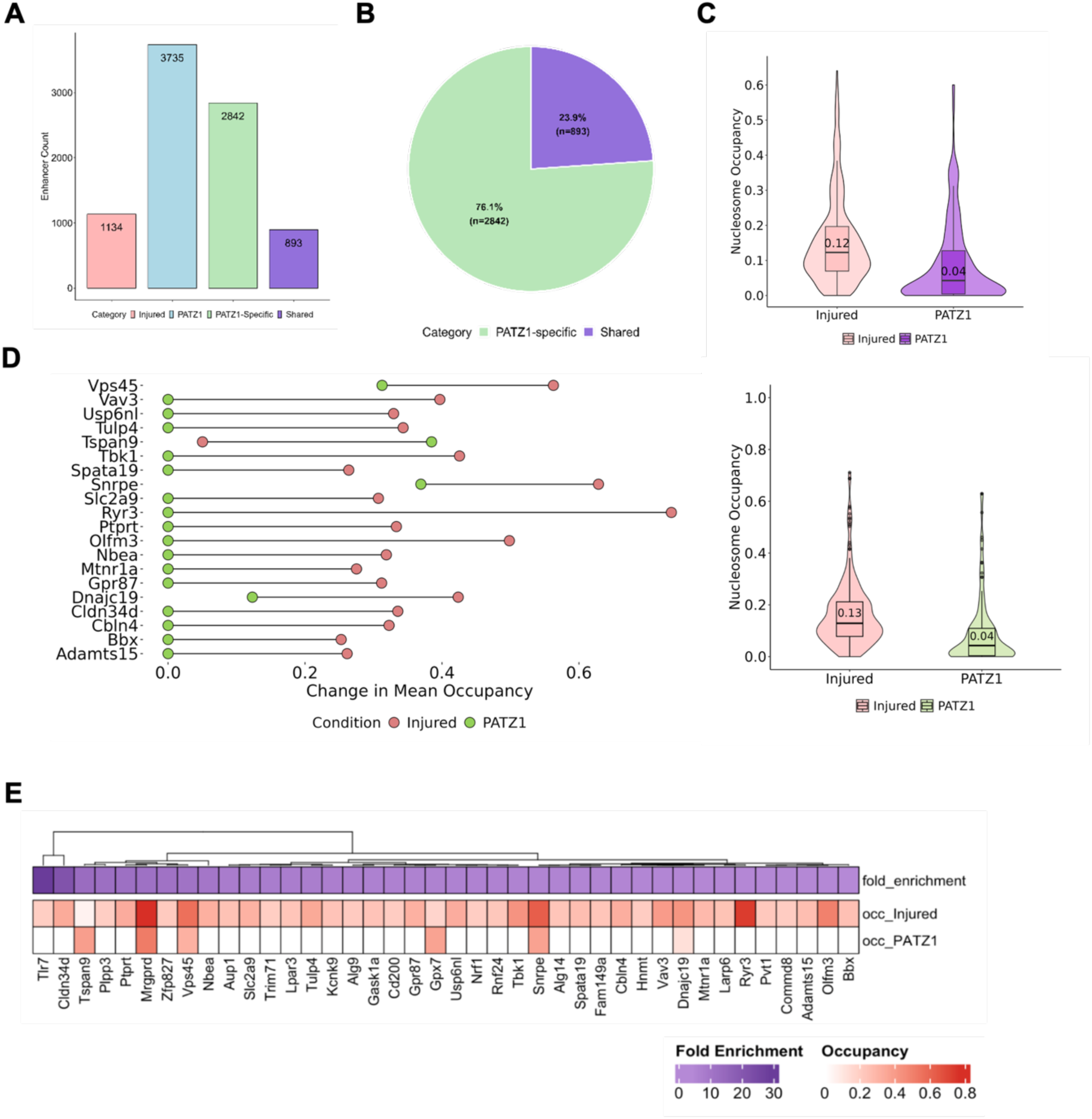
PATZ1 treatment of injured neurons reduces occupancy across shared and PATZ1-specific H3K27ac enhancer regions. **(A)** Bar chart showing the distribution of enhancer regions across 4 categories, wherein PATZ1 alone opens 2842 enhancer regions. **(B)** Pie chart highlighting the percentage of shared (purple) and PATZ1-specific (green) enhancer regions. **(C)** Violin boxplots comparing the nucleosome occupancy. The above panel shows the distribution in shared enhancer regions between injured (red) neurons and PATZ1-treated (violet) injured neurons. Median occupancy injured – 0.12; PATZ1 – 0.04. The lower panel shows the distribution in PATZ1-specific enhancer regions (injured – red; PATZ1 – green). Median occupancy injured – 0.13; PATZ1 – 0.04. **(D)** Lollipop plots showing the occupancy change across top 20 genes showing occupancy change in PATZ1-specific enhancer regions. **(E)** Heatmap comparing the fold enrichment scores from PATZ1-specific enhancer regions with the occupancy scores of injured and PATZ1-specific conditions.

Among enhancers with detectable nucleosome calls, the majority (76.1%, n=2842) were PATZ1-specific, while 23.9% (n=683) were shared between conditions (**Figure. 8B, Supplementary table S10**). This distribution indicates that PATZ1 preferentially acts on previously inaccessible enhancer regions rather than simply modifying existing accessible sites.

Nucleosome occupancy analysis at these enhancer categories revealed consistent depletion following PATZ1 treatment. At shared enhancer regions, median nucleosome occupancy decreased from 0.12 in injured neurons to 0.04 in PATZ1-treated neurons (**Figure. 8C, upper panel, Supplementary table S10**). PATZ1-specific enhancer regions showed a similar pattern, with median occupancy dropping from 0.13 to 0.04 between conditions (**Figure. 8C, lower panel, Supplementary table S10**). Both enhancer categories displayed substantially narrower occupancy distributions in the PATZ1 condition, reflecting more uniform nucleosome depletion across these regulatory elements.

Examination of individual genes associated with PATZ1-specific enhancer regions identified substantial occupancy changes at loci relevant to neuronal function and regeneration (**Figure. 8D, Supplementary table S10**). The top 20 genes showing occupancy reduction included *Vps45*, *Vav3*, *Usp6nl*, *Tulp4*, *Tspan9*, *Tbk1*, *Spata19*, *Snrpe*, *Slc2a9*, *Ryk*, *Ptprt*, *Olfm3*, *Nbea*, *Mtrn1a*, *Gpr87*, *Dnajc19*, *Cldn34d*, *Cbln4*, *Bbx*, and *Adamts15*. These genes showed high nucleosome occupancy in injured neurons that was markedly reduced following PATZ1 treatment, with occupancy changes ranging from approximately 0.2 to 0.6 units.

Heatmap analysis integrating fold enrichment scores with nucleosome occupancy across PATZ1-specific enhancer regions demonstrated a consistent relationship between enhancer activation and nucleosome depletion (**Figure. 8E, Supplementary table S10**). Genes including *Ttr7*, *Cldn34d*, *Tspan19*, *Plpp3*, *Ptprt*, *Mrgprd*, *Zfp827*, *Vps45*, *Nbea*, *Aup1*, *Slc2a9*, *Tmm71*, *Lpar3*, *Tulp4*, *Kcnk9*, *Alg9*, *Gask1a*, *Cd200*, *Gpr87*, *Gpx7*, *Usp6nl*, *Nrf1*, *Rnf24*, *Tbk1*, *Snrpe*, *Alg14*, *Spata19*, *Fam149a*, *Cbln4*, *Hmx3*, *Vav3*, *Mnjcl19*, *Mtrn1a*, *Larp6*, *Ryk3*, *Pvt1*, *Commd8*, *Adamts15*, *Olfm3*, and *Bbx* all showed high fold enrichment at PATZ1-specific enhancers coupled with elevated occupancy in injured neurons that was substantially reduced upon PATZ1 treatment.

These results demonstrate that PATZ1 not only remodels nucleosomes at existing accessible regions but also pioneers accessibility at thousands of previously closed enhancer elements, establishing an expanded regulatory landscape that may enable activation of regeneration-associated gene programs.

## Discussion

The inability of adult central nervous system (CNS) neurons to regenerate after injury remains one of the most significant challenges in restorative neuroscience^18^. While the field has historically focused on extrinsic inhibitory factors^5^, it is now clear that a major hurdle also lies within the neuron itself, specifically in the form of a growth-restrictive transcriptional^17,19–23^ and epigenetic landscape^6,8–12,24^. In this study, we investigate the molecular mechanisms through which the transcription factor PATZ1 orchestrates the regenerative potential of adult central nervous system (CNS) neurons. While our previous work identified PATZ1 as a potent driver of chromatin accessibility around growth genes^14^, the precise mechanisms underlying this transition remained elusive. Here, we demonstrate that PATZ1 promotes accessibility by actively modulating nucleosome occupancy. Our results reveal that PATZ1 induces widespread nucleosome eviction at critical regeneration-associated gene loci, significantly reducing both occupancy and precision to stabilize regulatory elements. This targeted remodeling is particularly evident at distal regulatory elements, where we observed a three-fold expansion of H3K27ac-marked active enhancers. By establishing that nucleosome eviction serves as the foundational step for enhancer activation, we provide a definitive mechanistic link between the physical clearing of the genomic landscape and the restoration of intrinsic growth capacity in injured CNS neurons.

Nucleosomes represent the fundamental repeating structural units of eukaryotic chromatin, with approximately 147 bp of DNA wrapped around a histone octamer core^25^.The positioning and occupancy of nucleosomes critically determine chromatin accessibility, as nucleosomes occluding DNA binding sites present formidable barriers to transcription factor binding and RNA polymerase machinery^26^. Transcriptionally active regulatory regions, including promoters and enhancers, are typically characterized by nucleosome-free or nucleosome-depleted regions that permit assembly of the transcriptional machinery^27^. The dynamic regulation of nucleosome positioning involves multiple mechanisms, including the action of ATP-dependent chromatin remodeling complexes that slide, eject, or exchange nucleosomes within chromatin. Recent high-resolution mapping approaches have revealed that nucleosomes at active regulatory regions exist in dynamically unwrapped states, with partial nucleosome disassembly facilitating transcription factor access while maintaining some degree of chromatin organization^16^. These unwrapped nucleosomal intermediates, often associated with SWI/SNF family remodelers, represent novel signatures of active chromatin that bridge nucleosome structure with transcriptional regulation. Additionally, cell type-specific differences in global nucleosome spacing have been documented, with neurons exhibiting shorter average spacing compared to glial cells, potentially reflecting their unique transcriptional requirements ^28^.

The failure of central nervous system axons to regenerate following injury, in contrast to the regenerative capacity of peripheral nervous system neurons, is increasingly understood to involve epigenetic mechanisms^29^. The current understanding of epigenetic regulation in the field of axon regeneration emphasizes a critical failure of CNS neurons to maintain or re-establish an open chromatin landscape following injury^13^. We previously established this connection by showing that during development, cortical neurons undergo a progressive “chromatin restriction” where sub-networks of genes associated with axon growth are systematically closed ^11,12^.While peripheral nervous system (PNS) neurons can remodel their epigenome following injury, marked by distinct signatures of H3K9ac and H3K27ac enrichment and increased accessibility, CNS neurons lack this intrinsic ability^10^. It is now appreciated that this lack of chromatin accessibility acts as a molecular “bottleneck,” preventing pro-regenerative transcription factors from accessing their target promoters in the adult brain or spinal cord ^13^.

Our work enhances this understanding by identifying nucleosome eviction as one of the molecular events that enable these broader epigenomic shifts. While previous studies have identified that the accumulation of histone marks or the complex 3D looping of DNA required for regeneration^10,24^, our results add a novel dimension and suggest that the physical removal of nucleosomes could be a critical prerequisite for these structural changes. By demonstrating that PATZ1-mediated eviction accompanies the expansion of active enhancers, we provide a high-resolution mechanism for how targeted remodeling is achieved in injured neurons.

While PATZ1-driven remodeling is robust, it remains unclear if the evicted nucleosomes eventually return or if the chromatin state is permanently altered. Future research should investigate the long-term stability of these nucleosome-depleted regions and whether they can be maintained long enough to support functional recovery in chronic injury models.

Additionally, while we focused on PATZ1, it is likely that other factors work in concert to coordinate nucleosome eviction with the 3D looping mechanisms involving cohesin that have been recently described in regenerating sensory neurons^24^. Understanding the hierarchy between nucleosome positioning, histone modification, and 3D folding will be essential for developing comprehensive regenerative therapies.

In summary, this study provides a new framework for understanding the structural barriers to CNS repair by linking PATZ1 activity to the physical eviction of nucleosomes. Our findings suggest that the regenerative potential of a neuron may not be permanently erased but may instead be buried under a nucleosomal architecture that can be strategically dismantled. Within the broad context of neurobiology, modulating nucleosome occupancy represents a promising avenue for epigenetic therapies aimed at “unlocking” the genome. By resetting the physical structure of chromatin, we can move closer to restoring functional connectivity and reversing the deficits caused by central nervous system injury.

## Methods

### Data acquisition and experimental groups

The single-nuclei ATAC (snATAC-seq) datasets used for nucleosome calling were derived from the motor cortex of adult mice subjected to three different experimental conditions. In each case, motor cortex tissue was isolated, and nuclei were extracted and sorted via Fluorescence-Activated Cell Sorting (FACS). The sorted nuclei were then processed following the 10X Genomics protocol (10x Genomics, CG000496 Chromium Next GEM Single Cell ATAC Reagent Kit v2). The study examined three biological conditions: a control group with uninjured tissue, an injury group with tissue collected seven days after a thoracic crush injury, and a treatment group where animals received AAV-PATZ1 for overexpression, followed by a thoracic crush injury. Detailed protocols for library preparation and sequencing are provided in ^14^, from which these datasets originated.

### Data Normalization and Nucleosome Positioning

Across all samples, an average sequencing depth of 10X coverage was achieved. To eliminate bias introduced by varying library sizes, all datasets were normalized by down-sampling to a uniform sequencing depth prior to downstream comparative analysis.

### Identification of broad peak regions

Chromatin accessibility peaks were identified from the snATAC-seq BAM files using MACS2 (version 2.2.9.1)^30^. To capture the wider genomic regions associated with nucleosome arrays, the callpeak function was employed with the --broad flag. The effective genome size was set for the mouse genome (mm10), and bed files were generated to facilitate signal visualization. These broad peak regions served as the spatial framework for the subsequent NucleoATAC pipeline^15^.

### Nucleosome calling and signal normalization

Nucleosome positions were identified using the NucleoATAC pipeline, which integrates chromatin accessibility data with sequence-specific biases. The analysis required three primary inputs: raw alignment files (BAM format), broad peak regions identified via MACS2 (BED format), and the mouse reference genome (mm10). The pipeline operates by cross-correlating a standard nucleosome V-plot with genomic fragment distributions. To ensure data integrity, the signal was normalized by subtracting background noise associated with Tn5 transposase insertion bias. The identification process was executed in two primary stages. First, nucleosome occupancy was calculated to generate occupancy scores and peak files. These results, along with the V-plot matrix, were then used to determine precise dyad coordinates. Additionally, nucleosome-free regions (NFRs) were computed by integrating the occupancy tracks with the final nucleosome calls. The resulting coordinate files provided the foundation for comparing nucleosome positioning across the three experimental conditions.

### Genomic annotation and feature distribution

To associate the identified nucleosome positions with biological functions, the NucleoATAC output files were annotated using the ChIPseeker package in R^31^. Nucleosome dyads and nucleosome-free regions (NFRs) were mapped to the mouse reference genome (mm10) using the annotatePeak function. This process assigned each genomic coordinate to specific features, including promoters, introns, exons, and intergenic regions. The distance from each nucleosome dyad to the nearest Transcription Start Site (TSS) was calculated to evaluate the distribution of chromatin accessibility relative to gene regulatory regions. For downstream visualization and comparative analysis between conditions, genomic feature distributions and TSS-distance profiles were generated using the ggplot2 package^32^.

### Statistical Analyses

Statistical analyses were performed using R version 4.3.2. Since the data for occupancy and fuzziness distributions did not meet the assumptions of normality, as assessed by the Shapiro-Wilk test and visual inspection of Q-Q plots, non-parametric methods were employed. The Kolmogorov-Smirnov test was utilized to compare the empirical distributions of the sample data against theoretical distributions. To evaluate differences in occupancy and fuzziness distributions across multiple groups, the Kruskal-Wallis H multiple comparison test adjusted with Bonferroni method was conducted. For significant results, post-hoc pairwise comparisons were performed using the Dunn test, with p-values adjusted for multiple comparisons using the Bonferroni correction to control the family-wise error rate.

The relationship between occupancy and fuzziness was quantified using Kendall’s tau (τ) correlation coefficient. This rank-based correlation was selected over Pearson’s *r* to better handle non-linear relationships, non-normal distributions and potential ties in the data. To interpret the magnitude of the observed differences, Cliff’s delta (δ) was calculated as a measure of effect size. Cliff’s δ was chosen as a robust non-parametric alternative to Cohen’s *d*, as it does not require assumptions regarding the shape or variance of the underlying distributions.

## Data availability

All genomics datasets generated in this study have been deposited in the NCBI Gene Expression Omnibus (GEO) under accession number **PRJNA1217518**. A detailed summary of datasets and corresponding sample information is provided in **Supplementary Table 1**. The metadata detailing the Supplementary Tables used for the results is provided in **Supplementary Table 11**. The code used for data processing and figure generation is available at https://github.com/mano2991/Targeted-chromatin-remodeller-PATZ1.

## Acknowledgements

We thank CSIR-CCMB for providing research facilities and support staff. We extend our thanks to the funding agencies Council of Scientific and Industrial Research (CSIR) (HCP53201), Department of Biotechnology (DBT) (BT/PR51467/MED/122/358/2024), and Science Engineering Research Board (SERB) (SRG00116), ANRF-SRG, CSIR Neuromission and BFIBiome.

## Author contributions

NK (Conceptualization, Data curation, Formal analysis, Investigation, Project administration, Validation, Visualization, Writing—review & editing), ASM (Data curation, Formal analysis, Investigation, Methodology, Project administration, Validation, Visualization, Writing—review & editing), MK (Data curation, Formal analysis, Software, Visualization, Writing—review & editing), DK (Methodology) and IV (Conceptualization, Data curation, Formal analysis, Funding acquisition, Investigation, Project administration, Resources, Supervision, Validation, Visualization, Writing).

